# Differential neurobehavioral responses to silica nanoparticles and temperature in a wild rhabditid nematode and *C. elegans*: toward an exposome map of phenotypes

**DOI:** 10.1101/2025.08.05.668664

**Authors:** A Limke, I Scharpf, A von Mikecz

## Abstract

Environmental factors shape organismal health through complex interactions known as the exposome. However, the interactions between chemical and non-chemical exposome factors remain largely unclear. Here, a dopaminergic reporter strain of *Caenorhabditis elegans* and a field isolated rhabditid nematode were used in a behavioral arena to assess natural variation in locomotory behavior in response to silica nanoparticles as a chemical exposome factor, and ambient temperature conditions (15 °C, 20 °C, and 25 °C) as non-chemical factors. Our results reveal that in the *C. elegans* strain lower temperature (15 °C) mitigates silica-induced locomotion deficits, while higher temperature (25 °C) exacerbates neurotoxicity, suggesting a temperature-dependent response. Notably, the wild rhabditid isolate showed distinct behavioral responses compared to the laboratory strain, highlighting the importance of species-specific ecological backgrounds in toxicological studies. This study provides a conceptual basis for integrating environmental factors and natural diversity into exposome research: Phenotype exposome maps, as presented here, broaden the identification of ecotoxicological hazards of nanomaterials across species.

**Author summary:** Incorporating wild rhabditid species with unknown genotypes into neurotoxicology research broadens the traditional model-organism framework by extending beyond *C. elegans* to encompass a wider range of naturally occurring neuronal and behavioral variation. This phenotype-driven approach enables the detection of both conserved and species-specific responses to neurotoxicants, independent of prior genomic knowledge. Observed differences in locomotor behavior across nematode species may reflect underlying diversity in neural architecture or compensatory mechanisms, offering insights into the functional plasticity of nervous systems. By adopting a comparative perspective grounded in the ecological context, this strategy enhances the environmental relevance of neurotoxicity studies and promotes more inclusive ecological risk assessment through the use of field isolated nematodes in controlled laboratory settings.

**Synopsis:** Wild nematodes and *C. elegans* differ in pollutant responses, highlighting the need for broader toxicology models.

**Highlights:**

- We report comparative testing of *Caenorhabditis elegans* and a wild rhabditid isolate in a behavioral arena.
- *C. elegans* and the wild isolate were assessed under multiple exposome conditions: silica nanoparticles and ambient temperatures (15–25 °C).
- Field isolated rhabditid nematodes showed distinct behavioral responses to chemical and temperature exposures compared to *C. elegans*.
- This study highlights the importance of including wild nematode isolates alongside *C. elegans* for ecological hazard identification of emerging pollutants like silica nanoparticles.
- We culture wild nematode isolates under standardized laboratory conditions to probe the limits of *C. elegans* as a toxicology model and enhance its real-world relevance.

## Introduction

The wild counterparts of laboratory model organisms such as *C. elegans*, *Drosophila*, and *Daphnia* live in environmental sinks of pollutants, including air, surface waters, sediments, and soil. These organisms serve as indicators for understanding the biological impact of complex environmental exposures. The exposome includes all chemical, biological, and physical exposures experienced by an organism over its lifetime (Gao, 2021; Zhang et al., 2021; Argentieri et al., 2025). Recent advances in environmental modeling make it possible to predict pollutant concentrations within environmental sinks (Mitrano et al., 2021; Keller et al., 2024), thereby enhancing our ability to map and quantify the external exposome more accurately.

To increase the ecological validity of molecular studies, scientists can perform molecular biology directly in the field or field isolate environmental samples, including wild invertebrates, for subsequent laboratory analysis (Heard, 2022; Perez DM et al., 2025). This emerging approach benefits from new technologies, such as automated behavioral platforms that assess motor-related phenotypes in *C. elegans* and its natural rhabditid relatives, enabling the classification and quantification of motility, including phenotypes associated with aging and neurodegeneration (Jung et al., 2015; Perni et al., 2018).

*C. elegans* is a widely used model organism in biomedical research, particularly for studying the molecular pathways involved in human disease. In the wild, it is a free-living soil organism that inhabits microbe-rich, organic environments typically found in the upper soil layers (Corsi et al., 2015). Due to its simplicity and genetic similarity to humans, the nematode serves as an effective system for understanding fundamental biological processes. Although *C. elegans* has a compact genome of about 100 million base pairs, it contains approximately 19,000 protein-coding genes, a number comparable to the roughly 20,000–21,000 protein-coding genes found in humans. About 60–80% of the nematode’s genes have human orthologs, often performing similar functions (Yoshimura et al., 2019). This genetic conservation extends to key biochemical pathways, such as insulin/IGF-1 signaling, which is also central to human aging and metabolism (Kaletta and Hengartner, 2006). These shared mechanisms make *C. elegans* a powerful model for studying conditions like neurodegeneration and metabolic disorders (von Mikecz, 2023). The *C. elegans* nervous system consists of exactly 302 neurons, which have been fully mapped at the synaptic level. It uses several conserved neurotransmitters, including acetylcholine, GABA, glutamate, dopamine, and serotonin. Transgenic reporter strains expressing fluorescent markers allow visualization of individual neurons that produce these neurotransmitters (Brewer et al., 2019; Santiago et al., 2022).

The nervous system of *C. elegans* is a primary target for pollutants, including certain nanoparticles, which can cause neurotoxicity characterized by neurodegeneration (Hua and Wang, 2023; Adedara et al., 2025). Damage to individual neurons during this neurotoxic process disrupts neurotransmission and neural circuits, leading to impaired neural function and altered behavior. The exposome, which encompasses all environmental exposures, both chemical and non-chemical, throughout an organism’s lifetime, provides a comprehensive framework for studying such neurotoxic effects (Gao, 2021; Gago-Ferrero et al., 2025). To investigate neurotoxicity within the context of the dynamic exposome, which includes multiple, interacting factors, we employed *C. elegans* on a behavioral platform to gain a more comprehensive understanding of real-world exposure scenarios.

Extending this approach, we applied the same testing paradigm to a field isolated rhabditid nematode to assess natural variation in response to environmental stressors. The wild isolate was used to assess natural variation in locomotory behavior upon exposure to silica nanoparticles as a chemical exposome factor and to ambient temperatures of 15° C, 20° C, and 25° C as non-chemical exposome factors. Observations of specific locomotion phenotypes such as bending, distance per bend, speed and area reveal that *C. elegans* reporter strains and wild rhabditid isolates respond differently to the combined environmental factors of nano silica and ambient temperature. The insights gained broaden our understanding of environmental stress responses and offer an initial contribution toward the development of an ecological exposome framework that integrates the One Health approach.

## Materials and Methods

### Strains and maintenance

The *C. elegans* reporter strain *egIs1 [dat-1p::*GFP] was obtained from the *Caenorhabditis* Genetic Center (CGC, University of Minnesota, Twin Cities, USA) and cultured on nematode growth medium (NGM) plates at 20 °C, using *Escherichia coli* OP50 as a food source. This reporter is commonly used to label dopaminergic neurons. In this strain, green fluorescent protein (GFP) is expressed under the control of the dat-1 promoter, which drives expression in the eight canonical dopaminergic neurons: four cephalic (CEP), two anterior deirid (ADE), and two posterior deirid (PDE) neurons (Nass et al., 2002). The integrated nature of the transgene ensures stable expression across generations, making it suitable for analyses of dopaminergic neuron function, and degeneration.

### Particles

Monodisperse silica nanoparticles (diameter: 50 nm) and BULK silica particles (diameter: 500 nm) were obtained from Kisker (Steinfurt, Germany). Fumed silica nanoparticles with nominal diameters of 12 nm (Aerosil® 200) and 20 nm (Aerosil® 90) were purchased from Evonik Industries (Essen, Germany). Stock suspensions (25 mg/mL in H_2_O) were prepared and sonicated using an ultrasonic homogenizer (Branson Sonifier 250-CE, G. Heinemann, Schwäbisch Gmünd, Germany). All particle dispersions were subsequently diluted in H_2_O as indicated. Particle characterization was performed by transmission electron microscopy (TEM) and dynamic light scattering (DLS), as previously described (Hemmerich and von Mikecz, 2013; Limke et al., 2024).

### Particle Exposures in Liquid Media

Nematodes were synchronized by isolating eggs via hypochlorite/NaOH treatment. At the L4 stage, animals were transferred to 96-well plates containing S-medium (pH 5.7) supplemented with 0.1 µg/mL Fungizone to prevent fungal contamination. For *Caenorhabditis elegans*, wells were seeded with freshly prepared *Escherichia coli* OP50 at a concentration of 12 mg/mL (Solis and Petrascheck, 2011). For field isolated rhabditid nematodes, a concentration of 15 mg/mL *E. coli* was used. To maintain developmental synchronization in *C. elegans* strain BZ555 (*egls1 [dat-1p::*GFP]), 5-fluoro-2′-deoxyuridine (FUdR) was added at a final concentration of 1.5 mM to inhibit reproduction. On day 1 of adulthood, all nematodes were either mock-exposed (H_2_O) or exposed to nano- or bulk-sized silica particles at the indicated concentrations. Liquid culture was conducted at 15 °C, 20 °C, or 25 °C.

### Field isolation of nematodes

Wild rhabditid nematodes were isolated from soil samples collected at two adjacent urban field sites near a heavily trafficked road in Düsseldorf, Germany (coordinates: 51°12’19.1"N, 6°47’12.4"E; 51°12’19.2"N, 6°47’10.6"E). Soil samples were transferred to nematode growth medium (NGM) agar plates seeded with *Escherichia coli* OP50 and incubated at 15 °C to allow nematodes to emerge, as described previously (Kim et al., 2014). Individual worms were isolated under a dissecting microscope and transferred to fresh plates for cultivation. Rhabditid nematodes were preliminarily identified and sorted based on morphological characteristics. For microscopic examination, nematodes were mounted on 5% agarose pads containing 10 μM sodium azide (NaN₃) and observed at room temperature using a stereomicroscope (SMZ18, Nikon Europe B.V., Amsterdam, Netherlands) equipped with a SHR Plan Apo 2× objective. Identification was based on key morphological features, including pharyngeal structures, vulva position, and intestinal pigmentation. (Sudhaus and Fitch, 2001; Lightfoot et al., 2016).

### Behavioral platform

Day 4 adult worms were transferred from 96-well microplates to 6-well agar plates overlaid with 2 mL of M9 buffer. After a 30-second adaption period to minimize stress-induced behavior, swimming was recorded using a Nikon SMZ18 stereomicroscope (Nikon Europe B.V., Amsterdam, The Netherlands) equipped with an SHR Plan Apo 1× objective and a DS- Qi2 digital camera. Videos were captured at 21 frames per second for 30 seconds using NIS Elements software (version 5.21.03, Nikon). Video analysis was performed using the Wide-Field Nematode Tracking Platform (WF-NTP), which incorporates a custom-written tracking algorithm capable of detecting individual nematodes based on their centroid positions. The software extracted key behavioral parameters, including swimming speed, body bend frequency, trajectory patterns, distance per bend, and morphological features. Worms were classified as paralyzed if they exhibited fewer than 0.5 body bends per minute and moved at less than 6 mm/min; such worms were marked as zero in the output Excel files. Analysis focused on behavioral and morphological parameters relevant to nematode locomotion. Body bends were quantified to assess rhythmic bending behavior reflective of neuromuscular function. Average and maximum speeds were measured to evaluate overall and peak locomotor activity, respectively. Movement efficiency was assessed by calculating distance per bend, which integrates propulsion and postural dynamics. In addition, body area was extracted as a morphological parameter. WF-NTP also generates color-coded locomotion tracks that visualize individual worm movement over time which facilitates the analysis of movement patterns under different conditions (Perni et al., 2018). The WF-NTP software is open-source and freely available at: https://github.com/impact27/WF_NTP (Koopman et al., 2020).

### Wide Field-of-View Nematode Tracking Platform (WF-NTP) - Parameters

In the WF-NTP platform, **average speed** refers to the mean swimming velocity of a worm, calculated as the average displacement per second across all tracked frames. In liquid environments, this parameter reflects overall swimming activity and is influenced by neuromuscular function or pharmacological treatment. **Maximal speed** captures the highest instantaneous velocity achieved during the observation period and can indicate brief bursts of rapid movement, such as escape responses or motor abnormalities. The **number of bends per 30 seconds** (30 s) quantifies the C-shaped body bends characteristic of swimming, offering a direct measure of swimming frequency. This metric is sensitive to changes in neuronal or muscular output. **Distance per bend** is defined as the total distance traveled divided by the number of bends in the same time window. It serves as an indicator of swimming efficiency, reflecting how effectively each undulatory movement contributes to forward propulsion.

### Single worm tracking (SWT)

In *C. elegans,* 21-146 worms (n) were analyzed per treatment in each experiment. For field isolated rhabditid nematodes, the range was 25-166 worms (n) per treatment per experiment. Variation in the number of analyzed worms was due to exclusion criteria set in WF-NTP, such as worms swimming out of the frame or collisions. All experiments were performed at least in triplicate. The total worm throughput (N) was 11,029 for *C. elegans* and 21,486 for field isolated rhabditid nematodes (Supplementary Table 1).

### Aldicarb assay

Aldicarb was used as a positive control in experiments with field isolated rhabditid nematodes. Aldicarb inhibits acetylcholinesterase, leading to the accumulation of acetylcholine in the synaptic cleft and ultimately causing paralysis in nematodes. A 100 mM stock solution was prepared by dissolving aldicarb in 70% ethanol and diluted in H_2_O to a final concentration of 0.1 mM for experiments. Worms were exposed to aldicarb in M9 buffer for 30 minutes prior to locomotion tracking (Mahoney et al., 2006).

### Data analysis and statistics

Following behavioral data extraction via WF-NTP, the raw output was further processed using a custom R script (R version 4.5.0). This script performs automated data structuring, calculation of descriptive statistics, generation of data visualizations (spider webs, box plots, scatter plots), and statistical analyses. Specific statistical tests are indicated in the corresponding figure legends and were selected based on data characteristics. Briefly, normality was assessed using the Shapiro–Wilk test, and homogeneity of variances was evaluated with Levene’s test. If the data met the assumptions of normality and homogeneity, one-way ANOVA followed by Tukey’s post hoc test was applied. For non-parametric or heterogeneous data, the Kruskal–Wallis ANOVA with Dunn’s post hoc test was used. *P* values < 0.05 were considered statistically significant. The custom R script was generated with support from an AI-driven programming assistant (ChatGPT, OpenAI) and its accuracy was verified by manually reproducing key data processing steps in Excel (Microsoft Corporation) and confirming statistical results using OriginPro 2022 (OriginLab Corporation).

## Results and Discussion

### Silica nanoparticles alter locomotion behavior in *C. elegans* strain *egIs1* [*dat-1p*::GFP]

In order to assess the impact of silica nanoparticles (NPs) on neuronal behavior in *C. elegans*, a semi-automated single-worm tracking platform was used to quantify locomotion phenotypes. Worms were cultivated in liquid culture on 96-well plates and exposed to different silica nanomaterials including (i) monodisperse NPs with a diameter of 50 nm, (ii) Aerosil 90 with a primary particle diameter of 20 nm or (iii) Aerosil 200 with a primary particle diameter of 12 nm (van Ommen et al., 2012; Albers et al., 2015). After transfer to 6-well plates, locomotion was recorded by video microscopy, visualized as individual worm thrashing tracks, and quantified using an open-source analysis pipeline tailored to our single-worm tracking setup. (Figure 1A).

**Figure 1.**
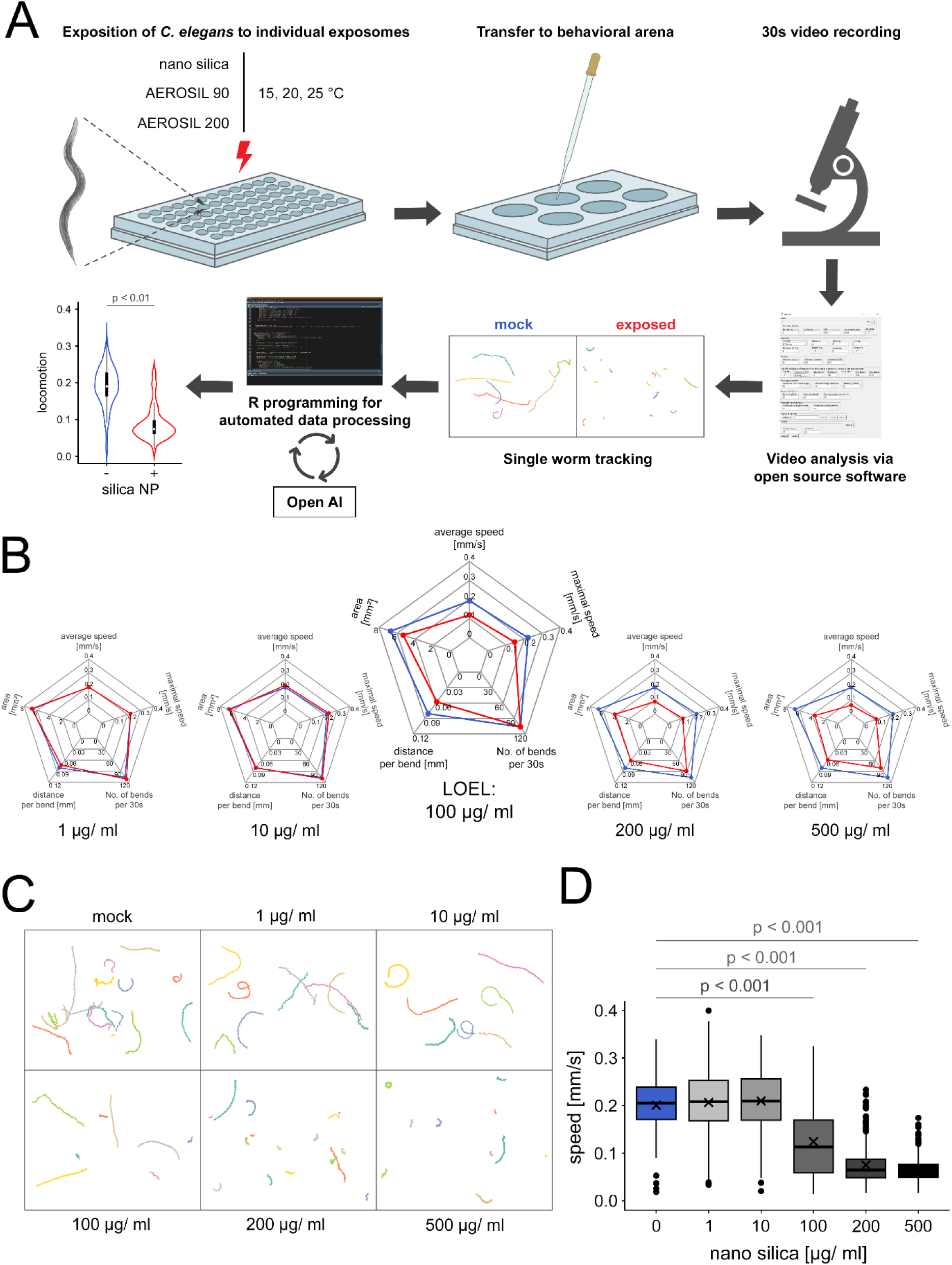
Quantification of locomotion in *C. elegans* exposed to silica nanoparticles using semi-automated single-worm tracking. (A) Schematic representation of the experimental workflow. Adult hermaphrodites of the reporter strain *egIs1* [*dat-1p*::GFP], which labels dopaminergic neurons, were exposed for 72 hours to increasing concentrations (1, 10, 100, 200, and 500 µg/mL) of three different silica nanomaterials: nano-silica (50 nm in diameter), Aerosil 90, or Aerosil 200. Controls were mock-exposed with H_2_O. All exposures were conducted in 96-well microtiter plates containing liquid medium and incubated at 15 °C, 20 °C, or 25 °C. After exposure, nematodes from individual wells were transferred to 6-well agar plates containing M9 buffer. Swimming behavior was recorded in each well for 30 s. Automatic video analysis was performed using the Wide Field-of-View Nematode Tracking Platform (WF-NTP) software to visualize individual swimming tracks of single worms and extract specific locomotion-related parameters. (Koopman et al., 2020). Unexposed controls show movement over long distances, whereas silica-exposed nematodes show reduced movement. The video analysis pipeline was extended with automated data processing using a custom R script developed via OpenAI-assisted prompting, enabling automated data organization, visualization and statistical analysis. (B) Representative spider plots show the means of several locomotion parameters — average speed, maximum speed, number of bends per 30 s, distance per bend, and area — after 72 hours of exposure to increasing concentrations of nano silica or H_2_O (control) at 20 °C. Data from the H_2_O control group are shown in blue, while data from nano-silica–exposed nematodes are shown in red. A concentration-dependent decline in locomotor function was observed, particularly in average and maximum speed, identifying these as the most sensitive locomotion parameters to silica nanoparticle exposure. The lowest observed effect level (LOEL) was determined to be 100 µg/mL. Based on this finding, speed was selected for detailed analysis in subsequent figures. (C) Representative swimming tracks of individual worms (multi-colored) under identical conditions are shown. The mock control group displays long tracks, indicating sustained locomotor activity. In contrast, exposure to increasing concentrations of nano-silica results in progressively shorter tracks, reflecting impaired mobility. The first noticeable reduction in swimming behavior is observed at 100 µg/mL, consistent with the identified LOEL. (D) Box plots represent the average speed of *C. elegans* after 72 hours of exposure to increasing concentrations of silica nanoparticles at 20°C. Data are presented as mean ± SD from four independent experiments, with n ≥ 26 worms per condition in each experiment. Statistical analysis was performed using Kruskal–Wallis ANOVA followed by Dunn’s post hoc test.

Measurements of different locomotion parameters showed that, with increasing concentrations of monodisperse silica NPs average speed and maximal speed were the most sensitive gait parameters, showing a significant reduction at concentrations of 100 μg/ml and above (Figure 1B). In contrast, the commonly used parameter ‘bends per 30 seconds’ was not as sensitive at the lowest observed effect level (LOEL) of 100 μg/ml. The parameters ‘distance per bend’, and ‘area’ showed intermediate sensitivity, but were less effective than speed as read outs (Figure 2B). In conclusion, a focused analysis of locomotion traits in a behavioral arena revealed that speed is a sensitive and reliable parameter for quantifying neuronal function following exposure to emerging pollutants such as nanoparticles.

**Figure 2.**
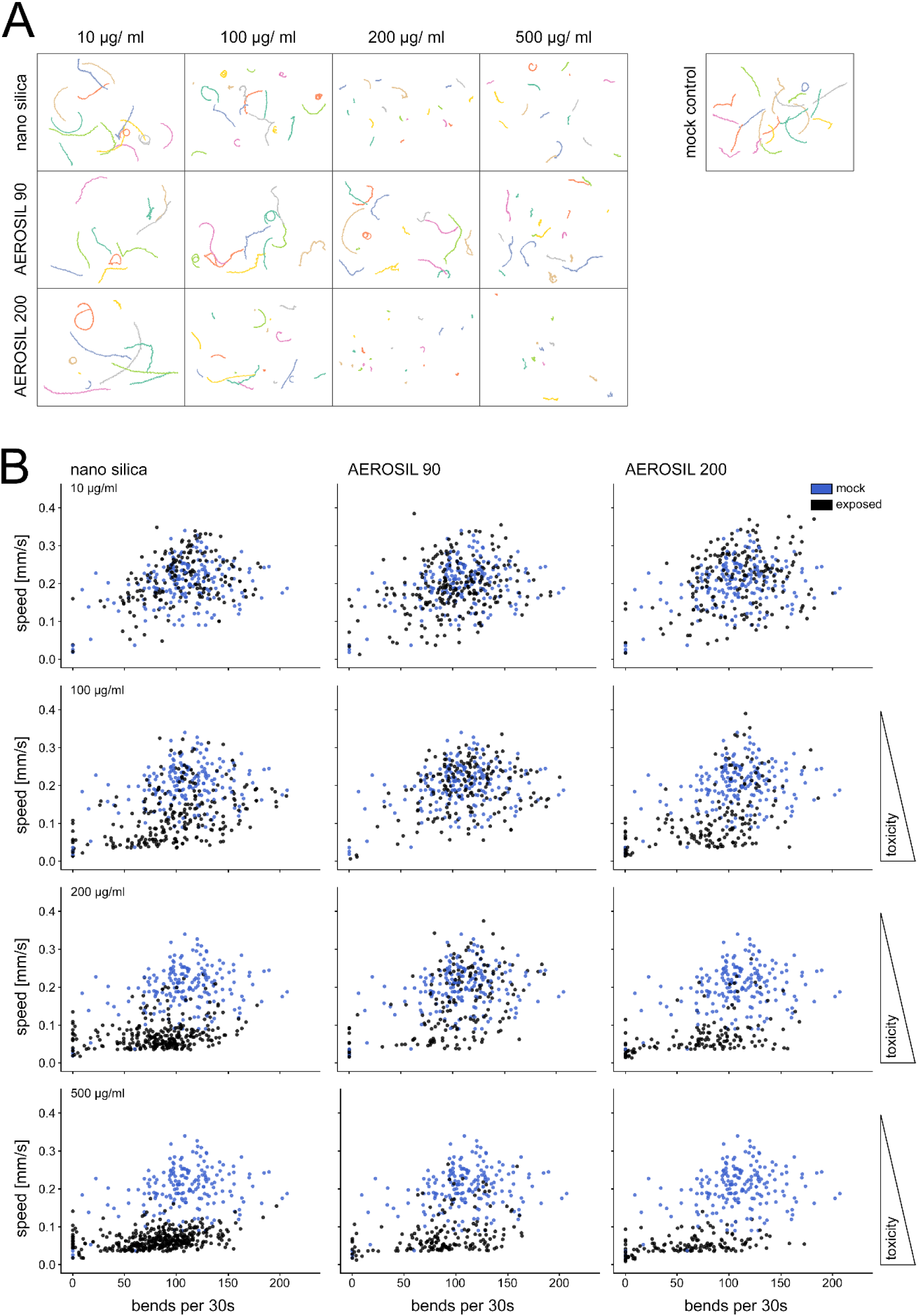
Differential effects of silica nanoparticles on locomotor phenotypes in *C. elegans* reporter strains. (A) Representative thrashing tracks are shown as tiled images of individual adult hermaphrodites from the reporter strain *egIs1* [*dat-1p*::GFP] (multi-colored), after 76 hours of exposure at 20°C to nano-silica, Aerosil 90, or Aerosil 200 at concentrations of 10, 100, 200, or 500 µg/mL (black), or to H_2_O as a mock control (blue). Worms exposed to increasing concentrations exhibit increasingly shorter tracks, indicating impaired motility. (B) Scatter plot analysis of *C. elegans* swimming behavior. Each point represents an individual worm, plotted by the number of body bends per 30 s (x-axis) and mean swimming speed (y-axis). The parameter ‘average speed’ is defined as the median velocity in millimeters per second. ‘Bends per 30 s’ indicates the number of bending movements observed within a 30 second recording. Data points are color-coded by experimental group (unexposed controls, blue; silica nanoparticle-exposed, black); control data is displayed identically across all scatter plots and serves as a common reference for comparing the effects of each nanoparticle exposure group. The plot reveals distinct locomotion phenotypes: animals with high bend counts and high speed exhibit coordinated swimming (upper right quadrant), whereas worms with high bend frequency but low speed suggest uncoordinated movement (lower right). Low bend and low speed values indicate reduced or paralyzed motor function (lower left). Increasing concentrations of silica nanoparticle exposure (10 – 500 μg/ml) shift the distribution of data points toward the lower quadrants, indicating a reduction of swimming speed. Body bend frequency remains largely unchanged, suggesting that while rhythmic motor output is maintained, neuromuscular coordination and locomotor efficiency are impaired. Data shown represent n ≥ 26 worms per condition from four independent experiments. Statistical analysis was performed using Kruskal–Wallis ANOVA followed by Dunn’s post hoc test.

Next, thrashing tracks of individual worms were visualized in response to a 72-hour exposure to increasing concentrations of monodisperse silica NPs with a diameter of 50 nm (Figure 1C). Differently colored tracks of individual reporter *C. elegans* show long, straight tracks in the mock control, and a concentration-dependent increase in individuals displaying shorter or circular tracks, particularly at concentrations of 100 μg/ml and above (Figure 1C). Even at high concentrations, track length ranged from long thrashing tracks to virtually no observable motion, indicating a highly individual response, with some worms showing resilience and others susceptibility (Figure 1C, bottom row). The quantification of the entire population of exposed worms confirmed the LOEL of silica NPs at 100 μg/ml concerning the parameter speed, with further reductions at higher concentrations of 200 and 500 μg/ml (Figure 1D).

Taken together, these results demonstrate that exposure to silica NPs induces a dose-dependent alteration in locomotion behavior in *C. elegans*, revealing distinct phenotypic variability among individuals. The observed range from normal movement to severe impairment highlights both resilience and susceptibility within the population, underscoring the importance of single-worm analysis for understanding nanomaterial neurotoxicity.

### Different silica nanomaterials differentially impact *C. elegans* locomotion

In order to compare locomotion behavior in response to different silica nanomaterials, individual thrashing tracks were visualized following a 72-hour exposure to increasing concentrations of monodisperse silica NPs (nano silica), Aerosil 90, or Aerosil 200 (Figure 2A). In the mock control, reporter *C. elegans* exhibited long, straight tracks, while higher nanoparticle concentrations resulted in an increasing number of individuals showing shorter or circular movement patterns. The LOEL concentrations were 100 μg/ml for nano silica and Aerosil 200, and 200 μg/ml for Aerosil 90 (Figure 2A). Extremely short tracks or non-movers were observed in *C. elegans* reporters exposed to Aerosil 200 at concentrations of 200 µg/ml or higher. Thus, among the tested materials, Aerosil 200 had the most pronounced impact on locomotion.

Next, the locomotion behavior induced by different silica nanomaterials was analyzed using two-dimensional scatter plots (Figure 2B). Data of unexposed worms (N=187; blue color) and *C. elegans* exposed to monodisperse nano silica (N=1173), Aerosil 90 (N=766), or Aerosil 200 (N=647) were plotted according to two motion parameters: bends per 30 s (x-axis) and speed (y-axis). At concentrations of 10 µg/ml, data points for unexposed and nanomaterial-exposed *C. elegans* are evenly distributed across all four quadrants. At concentrations of 100 µg/ml and above, data points from exposed individuals shift toward the two lower quadrants, indicating a reduction in speed while bending frequency remains largely unchanged. This pattern represents a specific movement gait and highlights how the scatter plot can capture detailed locomotor traits induced by silica nanoparticles. Confirming the thrashing track results, monodisperse silica NPs and Aerosil 200 had a greater impact on *C. elegans* locomotion parameters than Aerosil 90.

We conclude that among the tested silica nanomaterials, distinct effects on locomotor behavior emerged. Combined analysis of thrashing tracks and scatter plots revealed that these NPs primarily reduce speed while bending frequency remains largely unaffected, indicating specific alterations in movement gait. Reduction of the parameter speed specifically suggests respective neuromodulation by serotonergic neurotransmission as serotonergic *C. elegans* neurons NSM and ADF are likewise targets of silica NP exposure (Vidal-Gadea et al., 2011; Limke et al., 2024). The integration of complementary movement assays highlights specific alterations in behavioral patterns that offer insight into how NP properties modulate neuromuscular function in *C. elegans*. Here, behavioral phenotyping revealed a concentration-dependent decrease in neural function.

### Combined effects of silica NPs and ambient temperature on *C. elegans* locomotion

To compare locomotion behavior in response to silica nanomaterials under varying environmental temperature conditions, individual thrashing tracks were visualized following a 72-hour exposure to increasing concentrations of monodisperse silica nanoparticles (nano silica) at 15, 20, and 25 °C (Figure 3). Single-worm tracking showed normal movement patterns at 15 °C from 1 to 100 µg/ml. Tracks shortened at 200, and 500 µg/ml; however, no immobile worms were recorded (Figure 3A). At 20 °C, a similar pattern was observed, with exposure to 200 and 500 μg/ml nano silica resulting in either markedly reduced track lengths or punctate tracks, the latter indicative of a complete loss of locomotor activity (Figure 3C). Notably, this effect was even more pronounced at 25 °C, where a significant reduction in movement occurred already at a LOEL of 100 µg/ml (Figure 3E). Taken together, these results indicate that higher temperatures exacerbate nano silica-induced locomotion deficits in the *C. elegans* reporter strain *egIs1* [*dat-1p*::GFP].

**Figure 3.**
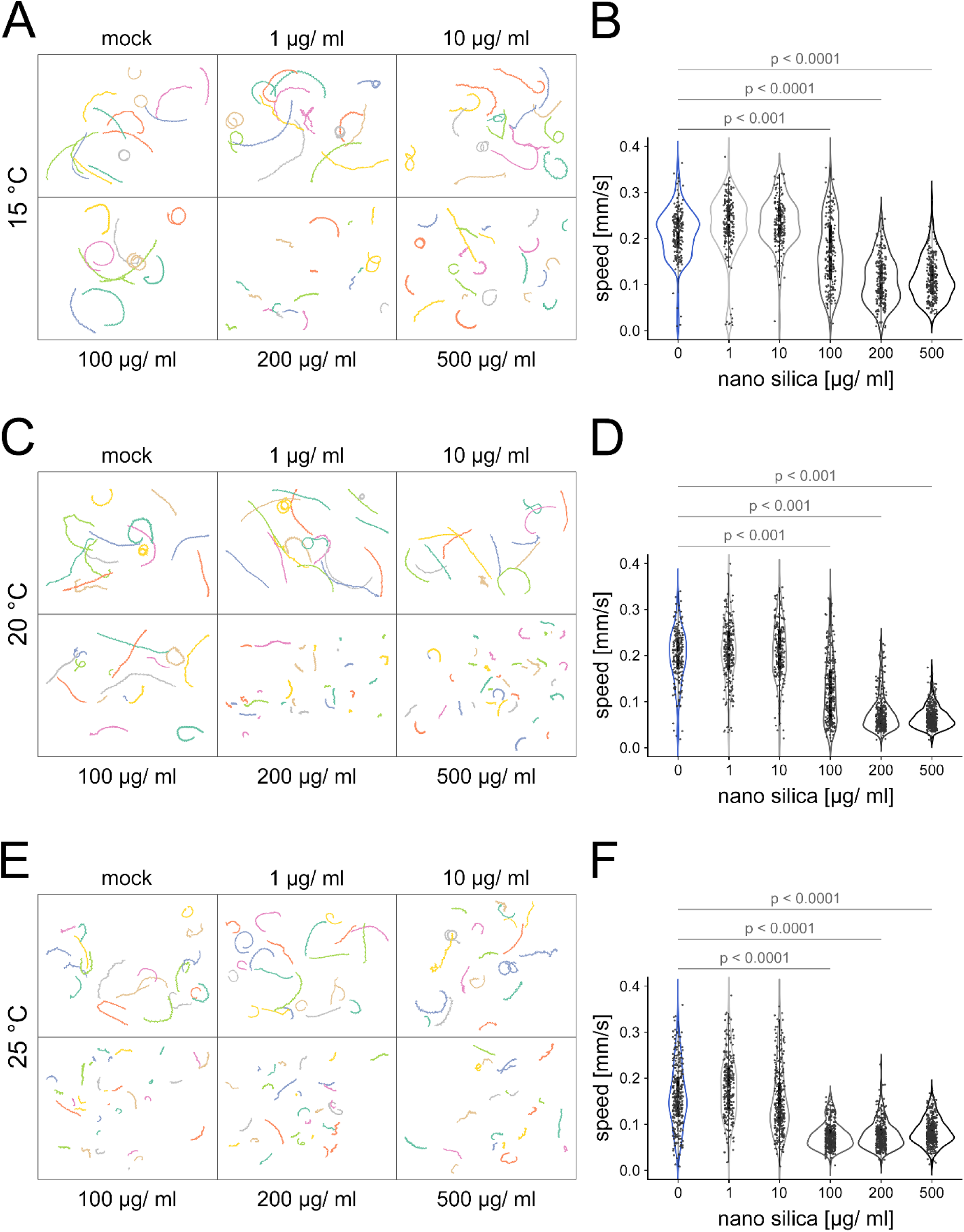
Temperature-dependent effects on locomotor phenotypes in *C. elegans* reporter strains exposed to silica nanoparticles. Representative thrashing tracks of individual adult hermaphrodites from the reporter strain *egIs1* [*dat-1p*::GFP] (multi-colored) are shown after 72 hours of exposure to increasing concentrations of silica nanoparticles (1, 10, 100, 200, 500 µg/mL) at (A) 15 °C, (C) 20 °C, or (E) 25 °C. Swimming activity is represented by ‘average speed’, defined as the median velocity of the worms in millimeters per second. The activity declined with increasing nanoparticle concentrations, as reflected by increasingly shorter swimming tracks. The reduction was more pronounced at higher temperatures, indicating temperature-enhanced neuromuscular defects. Tracks appearing as isolated points indicate a complete loss of locomotor activity. (B, D, F) Box plots represent the average speed of *C. elegans* under the same conditions at 15 °C (B), 20 °C (D), and 25 °C (F). Data are shown as mean ± SD from four independent experiments, with n ≥ 26 worms per condition in each experiment. Statistical analysis was performed using Kruskal–Wallis ANOVA followed by Dunn’s post hoc test.

Quantification of locomotion speed across the entire cohort confirmed a concentration-dependent decrease in movement, especially at 25 °C and concentrations of 100 µg/ml and above (Figure 3F). In conclusion, elevated temperatures intensify the locomotion-impairing effects of nano silica in *C. elegans*. These findings underscore the importance of considering temperature-related factors in nanomaterial toxicity assessments and demonstrate that single-worm tracking is a sensitive and effective method for detecting temperature-dependent impairments in locomotory behavior.

### Toward an exposome map of *C. elegans* locomotion

To investigate how environmental temperature and NP exposure influence *C. elegans* locomotion, we employed a wide field single-worm tracking platform (Perni et al., 2018). This system enabled the automated analysis of over 11,029 individual *C. elegans* hermaphrodites across a broad exposome-relevant parameter space.

Reporter strain *egIs1* [*dat-1p::*GFP] was exposed for 72-hours to three types of silica nanomaterials, monodisperse silica NPs (nano silica), Aerosil 90, and Aerosil 200, across five concentrations (1, 10, 100, 200, and 500 µg/mL) and at three ambient temperatures (15 °C, 20 °C, and 25 °C). Each condition was tested using 21 - 146 worms, and each experiment was independently repeated four times (Supplementary Figure 1; Supplementary Table 1).

Single-worm locomotion was recorded using automated time-lapse imaging at 21 frame/s for 30 s per worm, allowing for the precise extraction of quantitative behavioral features, including average speed, maximal speed, number of bends per 30s, distance per bend, area, and immobility events. This approach yielded over 55,145 discrete data points, providing a robust dataset to characterize dose- and temperature-dependent effects on locomotory behavior.

Together, this platform enabled detailed phenotyping of *C. elegans* locomotion in response to complex exposome variables and revealed the interactions between nanomaterial concentration and ambient temperature in driving locomotor impairment.

### Behavioral profiling of a wild rhabditid nematode isolate using automated single-worm tracking

Wild nematodes were isolated from soil samples collected in the field at 2 urban sites (Figure 4A). Soil was transferred to nematode growth medium (NGM) agar plates and incubated to allow nematodes to migrate out (Kim et al, 2014). Individual worms were isolated under a dissecting microscope and transferred to fresh plates for cultivation. Rhabditid nematodes were identified and manually sorted based on morphological characteristics (Figure 4C,D). The wild rhabditid isolate shows a strikingly similar appearance to an adult *C. elegans* hermaphrodite (Figure 4C). It has a typical vermiform and elongated body shape and measures approximately 1 mm in length. The body tapers at both ends and displays a smooth, cylindrical shape. Starting from the anterior, the wild isolate exhibits a head region with a visible pharynx. Under brightfield microscopy, the pharynx appears as a prominent structure with distinct compartments, namely the corpus, isthmus, and terminal bulb, the latter characterized by the presence of a grinder. (Figure 4D).

**Figure 4.**
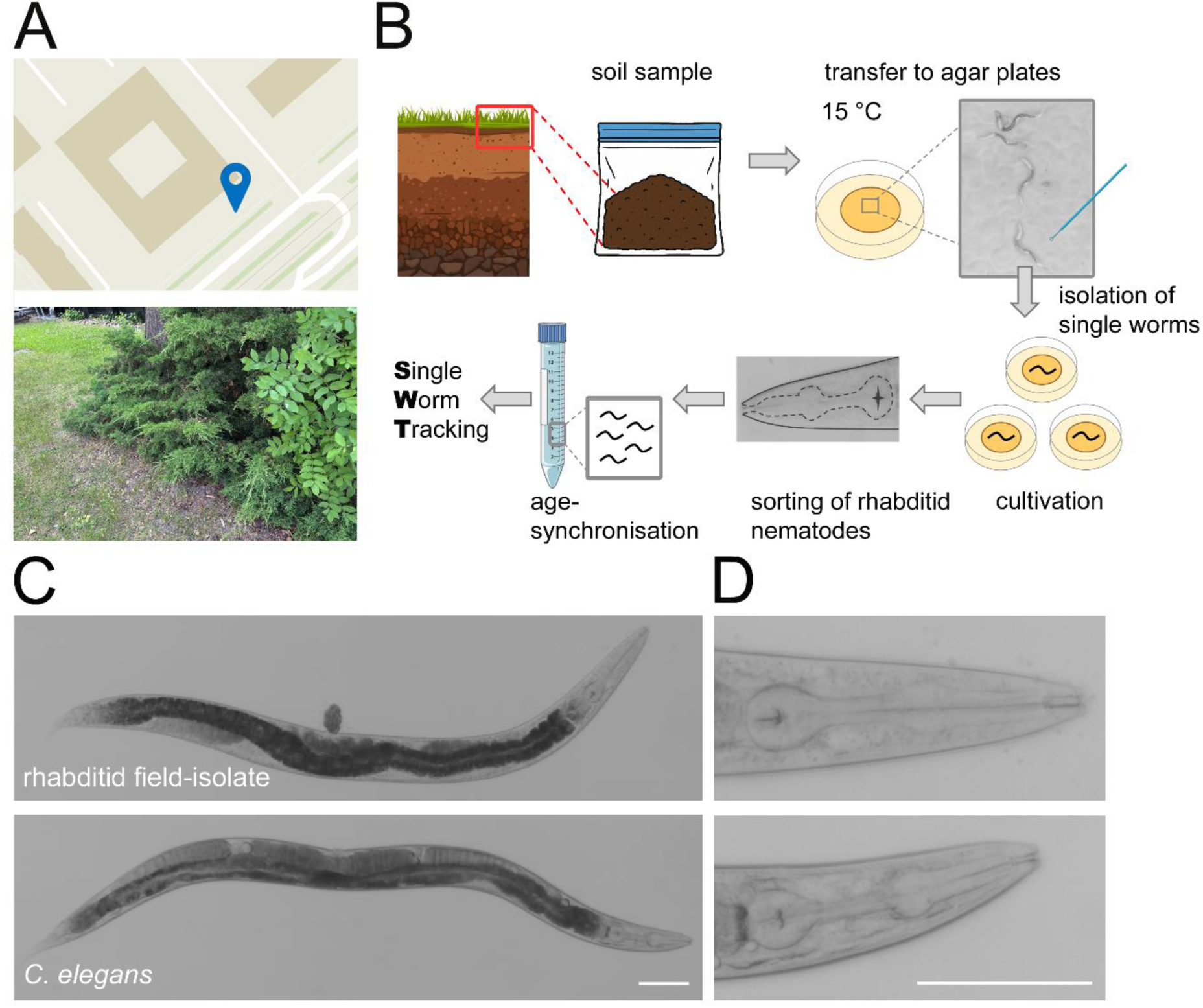
Field isolation and cultivation of wild rhabditid nematodes. (A) Schematic map of the field isolation site near a busy road in Duesseldorf, Germany (global coordinates: 51°12’19.1"N 6°47’12.4"E), and a photograph of the sampling site. (B) Soil was collected and applied to nematode-growth-medium (NGM) plates seeded with bacteria and incubated at 15 °C. Bacterivorous worms migrated from the soil onto the plates. Single worms were isolated, cultured, and identified as rhabditid nematodes based on morphological features (e.g., pharyngeal structure). Subsequently, age-synchronized worms were subjected to single worm tracking (SWT) analysis. (C) Micrograph of a representative field isolated rhabditid nematode compared to a *C. elegans* hermaphrodite (reporter strain *egIs1* [*dat-1p*::GFP]). (D) High-magnification micrograph showing the pharyngeal morphology of the wild rhabditid isolate and *C. elegans*, highlighting the terminal bulb with grinder, isthmus, and anterior bulb, as well as the characteristically long, straight stoma typical of the *Rhabditidae* family. Scale bars, 100 µm.

Posterior to the pharynx lies the intestinal tract, which extends as a wide, granular tube along most of the body. In the wild isolate as well as in *C. elegans* the intestine is surrounded by the reproductive system. Toward the posterior, the body narrows again, leading to the anus, which lies ventrally near the tail. Alongside the wild rhabditid isolates, other nematode species were also sampled from the field, including individuals resembling *Pristionchus* spp. These wild isolates exhibited distinct morphological differences from *C. elegans*, such as a toothed mouth, a pharynx lacking a grinder, and a shorter, stouter body with broader anterior region (Supplementary Figure 2).

Rhabditid field isolates were age-synchronized and subsequently used for behavioral assays on the single-worm tracking platform. Worms were either left unexposed or exposed to a concentration series of monodisperse silica NPs at 1, 10, 100, 200, and 500 µg/ml. Nanomaterial concentrations up to 100 µg/ml did not induce changes in locomotion phenotypes, whereas exposure to 200 µg/ml and higher resulted in a significant reduction in speed and area compared to unexposed controls (Figure 5A). Notably, visualization of single-worm tracks revealed that rhabditid field isolates generally moved at reduced speeds and followed shorter, less dynamic paths compared to *C. elegans* (Figure 5B). As a positive control, the acetylcholinesterase (AChE) inhibitor aldicarb was used, producing significant effects across all measured locomotion parameters, including speed, maximum speed, number of bends, distance per bend, and area. Consistent with this, single-worm tracks showed virtually no movement following aldicarb exposure (Figure 5B,C). Quantification of speed in the wild isolate population showed a reduction in locomotion following exposure to high concentrations of monodisperse silica NPs (200 and 500 µg/ml), although the effect was confined to a narrow activity range of approximately 0.05 and 0.1 mm/s (Figure 5C).

**Figure 5.**
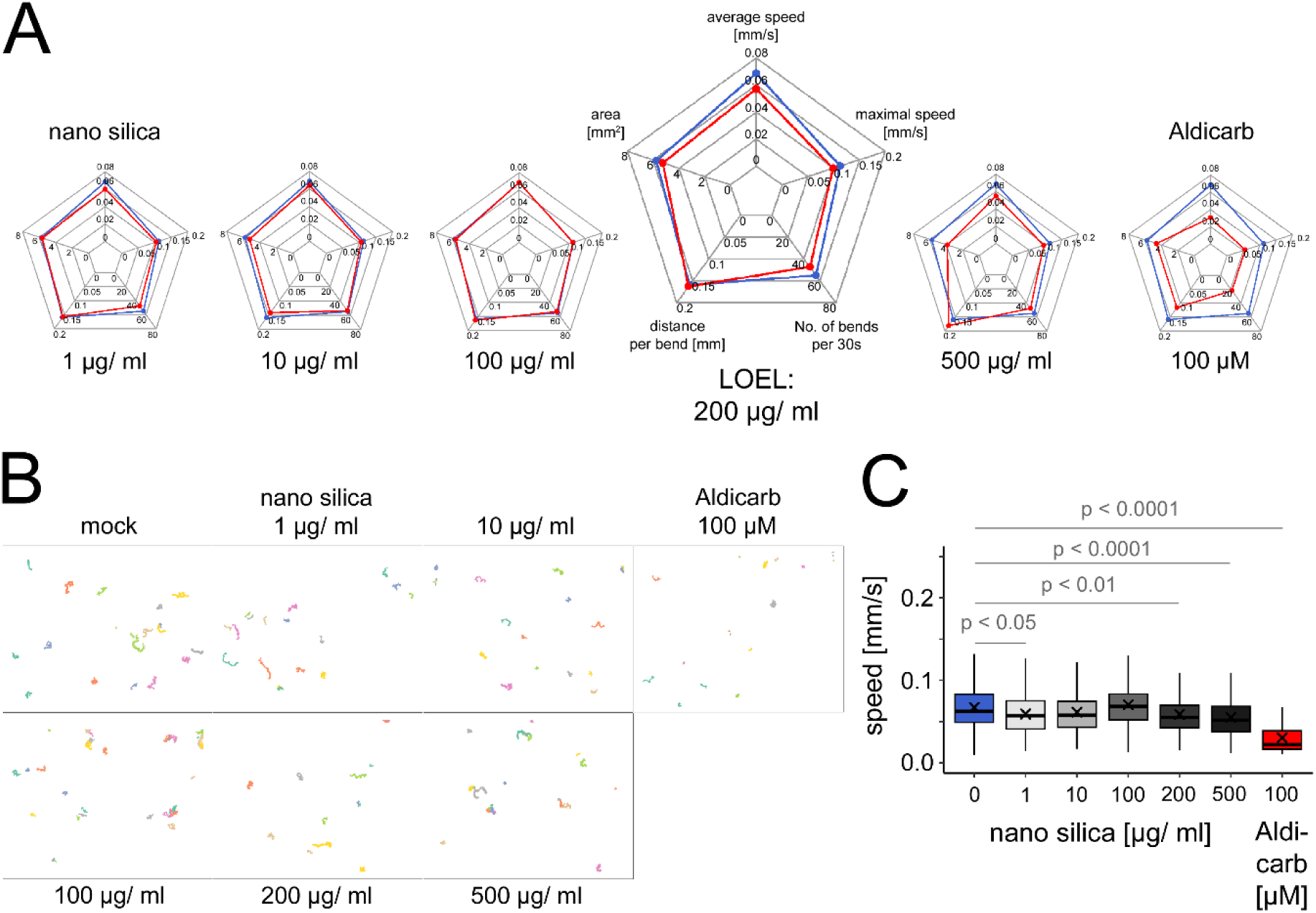
Single worm tracking with field isolated rhabditid nematodes. Field isolated rhabditid nematodes were exposed for 72 h to increasing concentrations of nano silica (1, 10, 100, 200 µg/mL), H_2_O (control) or 100 µM Aldicarb (positive control) at 20 °C. (A) Representative spider plots showing mean values of key locomotion parameters: average speed, maximum speed, number of bends per 30 s, distance per bend, and area. Control data are shown in blue; nano silica and Aldicarb-exposed groups are shown in red. Locomotor function declined in a concentration-dependent manner, with average speed, maximum speed, and number of bends showing the most pronounced reductions, indicating these as the most sensitive parameters. The lowest observed effect level (LOEL) was 200 µg/mL. Aldicarb reduced all locomotion parameters. (C) Representative 30-second swimming tracks of individual worms (multi-colored) under control, nano-silica, and Aldicarb conditions. Average speed was selected for detailed comparison with *C. elegans* in subsequent panels. (D) Box plots showing the average speed of worms exposed to increasing concentrations of silica nanoparticles or to Aldicarb. Data represent mean ± SD from three independent experiments, with n ≥ 37 worms per condition in each experiment. Statistical analysis was performed using Kruskal–Wallis ANOVA followed by Dunn’s post hoc test.

### Different silica nanomaterials differentially impact wild rhabditid isolates locomotion

In order to compare locomotion behavior in response to different silica nanomaterials in wild rhabditid isolates, individual thrashing tracks were visualized following a 72-hour exposure to increasing concentrations of monodisperse silica NPs (nano silica), Aerosil 90, or Aerosil 200 (Figure 6A). In the mock control, wild isolates exhibited short, zigzag or curvy tracks, while higher concentrations of monodisperse silica NPs resulted in an increasing number of individuals showing circular movement patterns or no movement. The LOEL concentration was 200 μg/ml for monodisperse silica NPs. In contrast, Aerosil 90 and 200, had no impact on thrashing tracks in comparison to the mock control (Figure 6A, middle and bottom rows). Thus, among the tested materials, monodisperse silica NPs had the most pronounced impact on locomotion.

**Figure 6.**
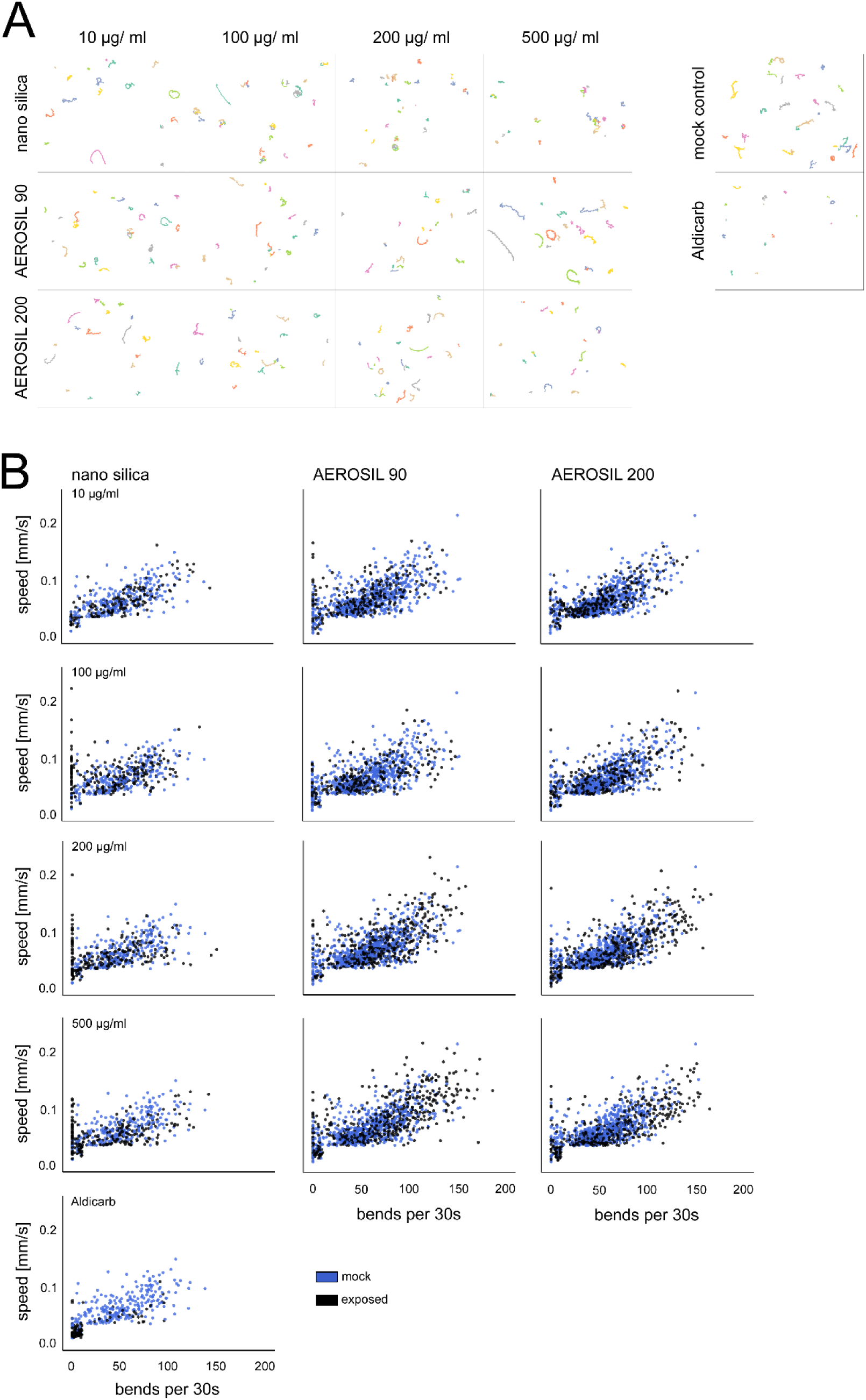
Multi-parameter analysis of locomotor phenotypes induced by silica nanoparticles in wild rhabditid nematodes. (A) Representative swimming tracks of individual adult rhabditid nematodes (multi-colored) after 72 h of exposure at 20 °C to nano-silica, Aerosil 90, or Aerosil 200 at concentrations of 10, 100, 200, or 500 µg/mL, to H_2_O (mock control), or to Aldicarb (positive control). Short tracks indicate reduced motility, while longer tracks reflect enhanced swimming. (B) Scatter plot of swimming behavior in individual worms, each represented by a point color-coded by treatment (controls: blue; silica-exposed: black). Data are plotted by body bends per 30 s (y-axis) and average swimming speed (x-axis). Increasing concentrations of nano silica and Aldicarb result in a shift toward the lower quadrants (lower speed, less bends), indicating impaired swimming and partial paralysis. In contrast, Aerosil 90 and Aerosil 200 cause a shift toward the upper quadrant (higher speed, more bends), suggesting enhanced locomotor activity. Data represent ≥32 worms per condition, from at least three independent experiments.

Next, locomotion behavior induced by different silica nanomaterials was analyzed using two-dimensional scatter plots (Figure 6B). Data of unexposed wild rhabditid isolates (N=1426); plotted in blue) and their counterparts exposed to monodisperse nano silica (N=1035), Aerosil 90 (N=1779), or Aerosil 200 (N=1887) were compared according to two motion parameters: bends per 30 s (x-axis) and speed (y-axis). At concentrations of 10 µg/ml, data points from both unexposed and nanomaterial-exposed wild rhabditid isolates were evenly distributed across all four quadrants. However, at concentrations of 100 µg/ml and above, a substantial fraction of data points from monodisperse silica NP-exposed individuals shifted toward the lower quadrants, and specifically along the y-axis, indicating a reduction or cessation of bending frequency.

In contrast, wild rhabditid isolates exposed to Aerosil 90 and Aerosil 200 displayed locomotion patterns distributed across all four quadrants. Notably, a subset shifted to the upper right quadrant, suggesting increased speed and thrashing rates. Another subset of wild isolates showed a concentration-dependent accumulation in the lower quadrant along the y-axis following exposure to all three particle types, indicating reduced speed and bending frequency.

Taken together wild rhabditid isolates exhibited a considerable diversity of movement patterns in response to the different silica nanomaterials. At the population level, opposing effects neutralized each other, resulting in an overall unchanged response to the pollutants. In contrast to the model organism *C. elegans* monodisperse silica NPs and Aerosil 200 had the strongest negative impact on locomotion behavior, whereas Aerosil 90 had little or even a stimulatory effect (Supplementary Figure 1).

A stimulatory effect was also observed following exposure of wild rhabditid isolates to monodisperse silica NPs at concentrations from 1 to 500 μg/ml at an ambient temperature of 15 °C (Supplementary Figure 1). Single-worm tracking revealed increased locomotor activity and longer movement tracks in individual worms (Supplementary Figure 3A). Consistent with these findings, quantification of locomotion speed across the entire cohort revealed a significant increase at both low and high concentrations of silica NPs (Supplementary Figure 3B). Notably, this represents the first reported observation of a stimulatory effect induced by silica nanomaterials, suggesting that field isolated wild nematodes may exhibit distinct responses to this chemical exposome factor. When we exposed the field isolates to silica particles with a diameter of 200 nm (BULK silica) a stimulatory effect was specifically observable at 15 °C (Supplementary Figure 4), suggesting that the material elicited the effect irrespective of particle size. Actually, this is likewise in contrast to previous results in *C. elegans* strains where all neurotoxic effects, including neural blebbing, bag of worms and intestinal ballooning were nano-specific and not inducible by bulk silica particles with diameters ≥ 200 nm (von Mikecz, 2023; Limke et al., 2024).

To further investigate the influence of the non-chemical exposome factor ambient temperature, the experiments were repeated at 20 °C and 25 °C. At these elevated temperatures, exposure to monodisperse silica nanoparticles led to a reduction in locomotion speed at concentrations ≥200 µg/ml at 20 °C and at 500 µg/ml at 25 °C, while lower concentrations had no observable effect (Supplementary Figure 3C-F).

These findings suggest that the behavioral response of nematodes to silica nanomaterial exposure is both temperature-dependent and species-specific. While wild rhabditid isolates exhibited a stimulatory locomotor response at 15 °C, this effect was reversed at higher temperatures (20 °C and 25 °C), where reduced speed was observed at elevated NP concentrations. In contrast, standard laboratory *C. elegans* strains did not display a comparable stimulatory effect, highlighting potential differences in environmental sensitivity or adaptive physiology between wild and domesticated nematode populations. These results underscore the importance of considering both non-chemical exposome factors, such as temperature, and the field origin of the test organisms when assessing the ecotoxicological impact of nanomaterials.

### Tracking behavioral responses to exposome factors in *C. elegans* and a wild rhabditid field isolate

To compare locomotion behavior in response to different silica nanomaterials, individual thrashing tracks were visualized in the *C. elegans* strain *egIs1* [*dat-1p*::GFP] and a wild rhabditid field isolate following 72-hour exposure to monodisperse silica nanoparticles (nano-silica), Aerosil 90, or Aerosil 200 (Figure 7A, top row). In the mock control, reporter *C. elegans* exhibited long, straight tracks, indicating wide-ranging movement. Exposure to monodisperse silica NPs and Aerosil 200 impaired locomotion, resulting in shorter, curved tracks. Aerosil 90 also reduced locomotion, but to a lesser extent.

**Figure 7.**
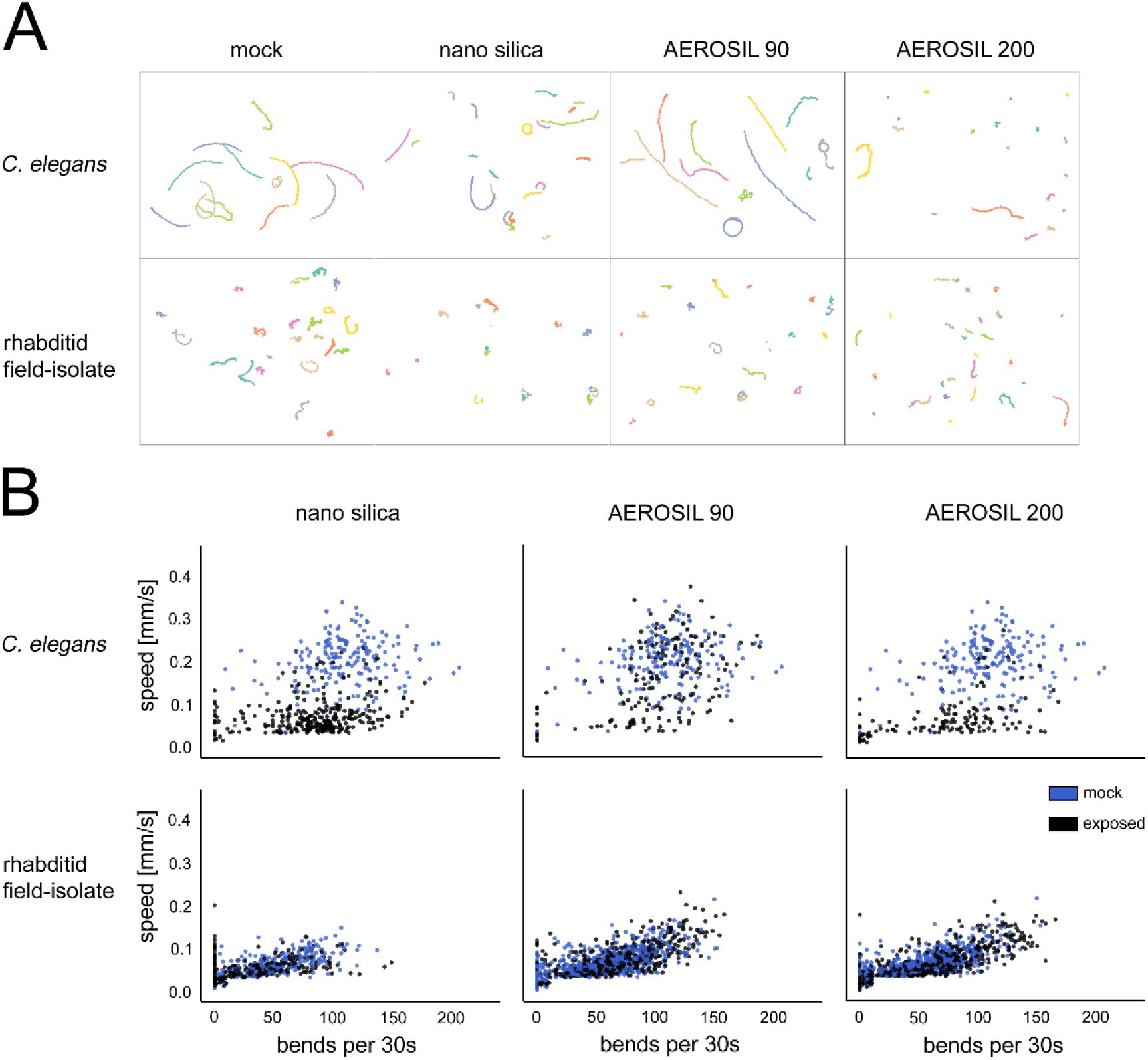
Comparison of single worm tracking in *C. elegans* versus field isolated wild rhabditid nematodes. (A) Comparison of individual swimming tracks (multi-colored) between *C. elegans* and field isolated rhabditid nematodes shows that *C. elegans* exhibits longer tracks, indicative of greater intrinsic locomotor activity under unexposed control conditions. Exposure to the LOEL (200 µg/mL) of monodisperse silica nanoparticles, Aerosil 90, or Aerosil 200 for 72 h at 20 °C results in visibly shorter swimming tracks in *C. elegans*, indicating reduced locomotion. In contrast, exposed field isolated rhabditid nematodes show no clearly discernible changes in swimming track patterns. (B) The scatter plot shows the swimming behavior of *C. elegans* and field isolated rhabditid nematodes after 72 h of exposure to monodisperse silica nanoparticles, Aerosil 90, or Aerosil 200. Average swimming speed is plotted against the number of body bends per 30 s. Each dot represents a single worm and is color-coded by treatment group (mock: blue; exposed: black). In *C. elegans*, mock- and particle-exposed groups form distinct clusters, indicating clearly altered locomotion phenotypes. Silica particle exposure shifts data point distributions toward the lower quadrants, indicating reduced swimming speed and bending frequency. In contrast, unexposed and exposed field isolated nematodes show substantial overlap in their distributions, suggesting minimal effects on locomotion. Nonetheless, distinct clustering patterns are detectable, and silica nanoparticles induce opposing effects on swimming behavior of the wild isolates. Exposure to nano-silica shifts data points toward the lower left quadrant, indicative of reduced locomotion or partial paralysis. In contrast, exposure to Aerosil 90 and Aerosil 200 shifts distributions toward the upper right quadrant, suggesting enhanced locomotor activity. Data represent ≥26 worms per condition from at least three independent experiments.

In contrast, the wild rhabditid isolates exhibited less wide-ranging movement in the unexposed control group compared to their *C. elegans* counterparts. Within this more restricted range, monodisperse silica NPs further reduced locomotory activity, whereas Aerosil 90 and 200 did not significantly affect locomotion. These findings indicate that monodisperse silica nanoparticles at a concentration of 200 μg/mL exerted the most pronounced effect on the wild isolates.

Next, the locomotion behavior induced by different silica nanomaterials was analyzed using two-dimensional scatter plots (Figure 7B). Data of unexposed *C. elegans* (top row, blue color) and *C. elegans* exposed to monodisperse nano silica, Aerosil 90, or Aerosil 200 were plotted according to two motion parameters: bends per 30 s (x-axis) and speed (y-axis). Data points for unexposed *C. elegans* are evenly distributed across all four quadrants. A similar pattern was observed in *C. elegans* exposed to Aerosil 90. However, data points from *C. elegans* exposed to monodisperse silica NPs and Aerosil 200 shift toward the two lower quadrants, indicating a reduction in speed while bending frequency remains largely unchanged.

Scatter plots of wild rhabditid field isolates show a different pattern. Unexposed as well as groups exposed to the different silica nanomaterials show movement patterns with a generally shorter range that distribute in the lower two quadrants. However, monodisperse silica NPs induce a reduction of both speed and bends per 30 s, indicating a significant reduction of locomotion activity in wild isolates.

These results reveal distinct locomotion responses to silica nanomaterials in *C. elegans* and wild rhabditid isolates. In *C. elegans*, exposure to monodisperse silica NPs and Aerosil 200 led to reduced speed without affecting bending frequency, while Aerosil 90 had minimal impact. In contrast, wild isolates showed consistently lower overall locomotion and responded to monodisperse silica NPs with a marked reduction in both speed and bending frequency. These findings suggest species-specific sensitivity, and the reasons for this variation remain to be explored.

Our use of field-collected rhabditid nematodes alongside the *C. elegans* laboratory reporter strain *egIs1* [*dat-1p*::GFP] aligns with the emerging “take the field to the lab” paradigm (von Mikecz & Scharf, 2022; Bean et al., 2024). This strategy enhances environmental realism by integrating organisms with natural ecological exposure histories into controlled laboratory settings. It preserves essential features such as local environmental imprint, while allowing for mechanistic investigation via standardized assays.

Visualization in an exposome map (Supplementary Figure 1) reveals pronounced differences in stress resilience between *C. elegans* and the wild rhabditid isolate. Under silica NP exposure, *C. elegans* shows reduced locomotion, whereas field-collected isolates maintain, or even increase, movement. This locomotor response is both temperature-dependent and species-specific: wild isolates exhibit a stimulatory effect at 15 °C, which diminishes or reverses at 20 °C and 25 °C. In contrast, *C. elegans* shows no comparable increase under any condition tested. These inter-species differences likely reflect distinct ecological exposure histories, indicating that environmental origin modulates nanomaterial sensitivity. Notably, this enhanced stress resilience in wild isolates may correlate with greater longevity, as preliminary results show significantly longer life spans compared to the laboratory strain (Limke and von Mikecz, unpublished).

Traditional nematode research, especially in molecular biology and toxicology, has relied on highly domesticated strains like N2, which thrive under uniform lab conditions (Brenner, 1974). While useful for reproducibility, these strains lack the genetic and phenotypic diversity present in wild populations. Field-collected nematodes provide an ecologically informed model system, enabling exploration of how natural environmental exposures interact with genetic background to shape responses to stressors such as nanoparticles and temperature variation.

Our findings highlight the value of incorporating wild isolates into exposome research. The differential behavioral responses between laboratory and field-derived strains emphasize the role of natural variation in shaping toxicological outcomes (Romero-Blanco & Alonso, 2022). As high-resolution tools like single-worm proteomics, transcriptomics, and barcoding become more accessible (Bensaddek et al., 2016; Tan et al., 2024; Zhu et al., 2024), the capacity to interrogate wild nematodes at molecular depth will expand, offering mechanistic insight grounded in ecological context.

Although still underutilized in molecular toxicology, this approach holds strong promise for advancing environmental health research. It supports a more integrated understanding of organismal responses to complex stressors and contributes to aligning toxicological models with real-world conditions, central to exposome science and the One Health framework. Nematode-based studies on pollutant-induced neurotoxicity can uncover conserved genes and pathways, which may map to human orthologs. These findings can inform population-level analyses, such as UK Biobank studies, by enabling gene–environment interaction mapping for pollutants and neurological outcomes (Argentieri et al., 2025). Moving forward, priorities include protocol standardization, population-level validation, and building a methodological foundation to support broader adoption of field-derived models. Further research is needed to elucidate how environmental heterogeneity and biological diversity shape responses to both emerging and legacy pollutants.

## Acknowledgements

Strains were provided by the *Caenorhabditis* Genetics Center (CGC), which is funded by the NIH. We are also grateful to members of the von Mikecz lab for their support, discussions, and guidance throughout this work. Additional thanks go to funding by the Deutsche Forschungsgemeinschaft, Grant MI 486/10-1 (rewarded to AvM) and the IUF. The IUF is funded by the federal and state governments - the Ministry of Culture and Science of North Rhine-Westphalia (MKW) and the Federal Ministry of Education and Research (BMBF).

## Conflict of Interest Statement

The authors declare that they have no competing interests.

## Author contributions

AvM: Conceptualization, Supervision, Writing of original draft, Resources, Funding acquisition. AL: Investigation, Methodology, Formal analysis, Data curation, Visualization, Writing. IS: Investigation, Methodology, Formal analysis, Data curation, Visualization, Writing. All authors reviewed and approved the final manuscript.

## Supplementary Material

**Supplementary Figure 1.**
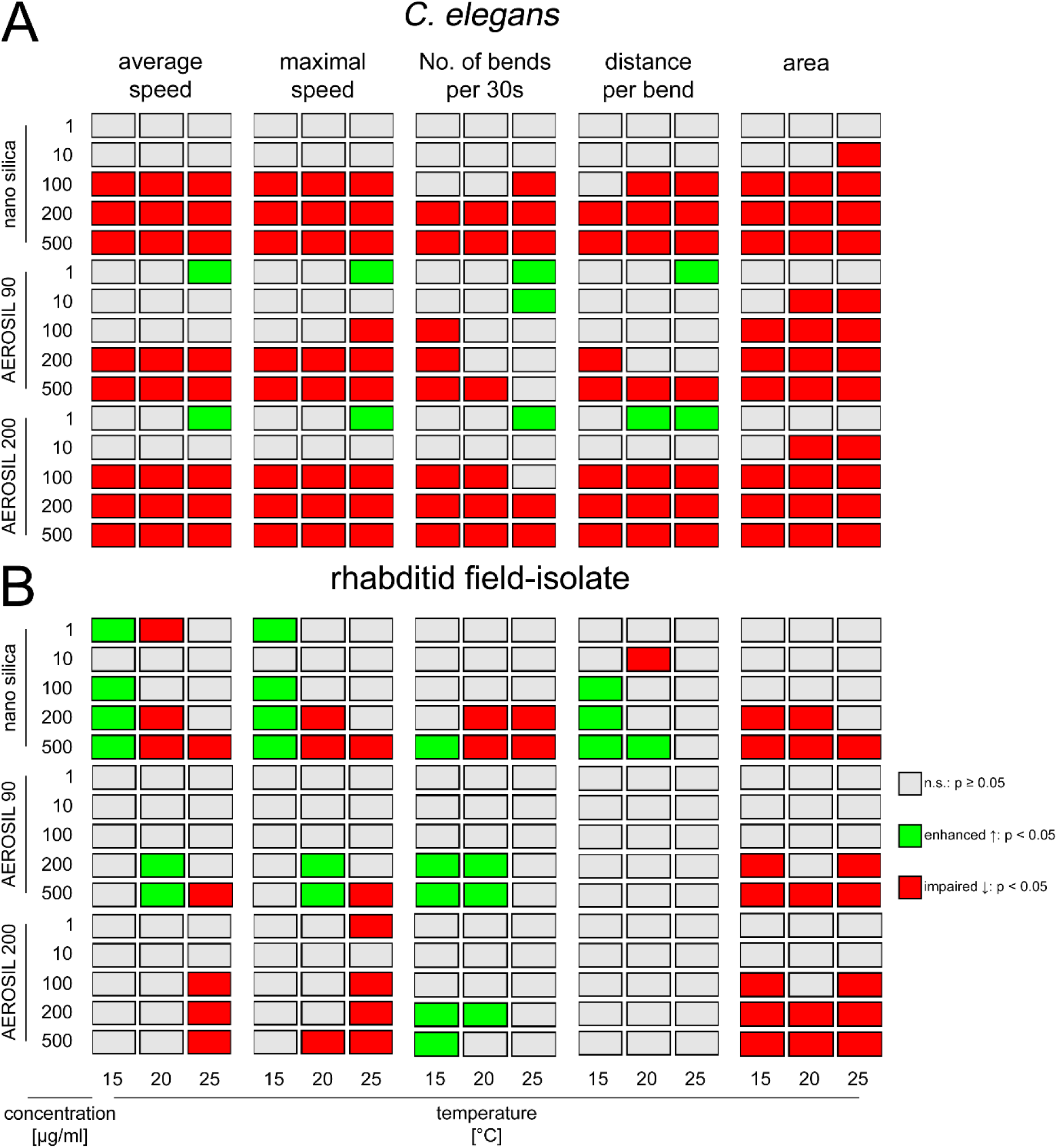
Exposome map – Nematode locomotion behavior was recorded and quantified using a wide-field single-worm tracking platform. (A) *C. elegans* strain *egIs1* [*dat-1p*::GFP] was exposed to increasing concentrations of monodisperse silica nanoparticles (nano silica), Aerosil 90, or Aerosil 200 at ambient temperatures ranging from 15 °C to 25 °C. (B) Same experiments with a field isolated rhabditid nematode. The exposome map provides a comprehensive overview of silica nanoparticle-induced neuromuscular locomotion behavior. It integrates multiple exposome factors: three particle types (monodisperse silica nanoparticles with a diameter of 50 nm, Aerosil 90, Aerosil 200), increasing concentrations (1–500 µg/mL), and ambient temperatures of 15, 20, and 25 °C. The map summarizes statistical significance (p-values) for five locomotion parameters: average speed, maximal speed, number of bends per 30 s, distance per bend, and area, across all experimental conditions. The parameter ‘average speed’ is defined as the median velocity in millimeters per second, while ‘maximum speed’ is estimated using the 90th percentile of speed values to represent peak velocities. ‘Bends per 30 s’ indicates the number of bending movements observed within a 30 second recording. The parameter ‘distance per bend’ quantifies the distance travelled during a single bending motion, expressed in millimeters. ‘Area’ provides an estimate of the worm body area, measured in square millimeters. Color coding indicates phenotypic outcomes and significance: light gray indicates no statistically significant change compared to unexposed controls (p ≥ 0.05), green indicates a significantly enhanced locomotion phenotype (p < 0.05), and red represents a significantly impaired locomotion phenotype (p < 0.05). The visualization highlights condition-specific and exposure-dependent behavioral effects, revealing particle- and temperature-specific neurotoxic profiles. Each condition included a minimum of n ≥ 21 worms per experiment. Experiments were conducted with at least three replicates. Statistical analysis was performed using Kruskal–Wallis ANOVA followed by Dunn’s post hoc test.

**Supplementary Figure 2.**
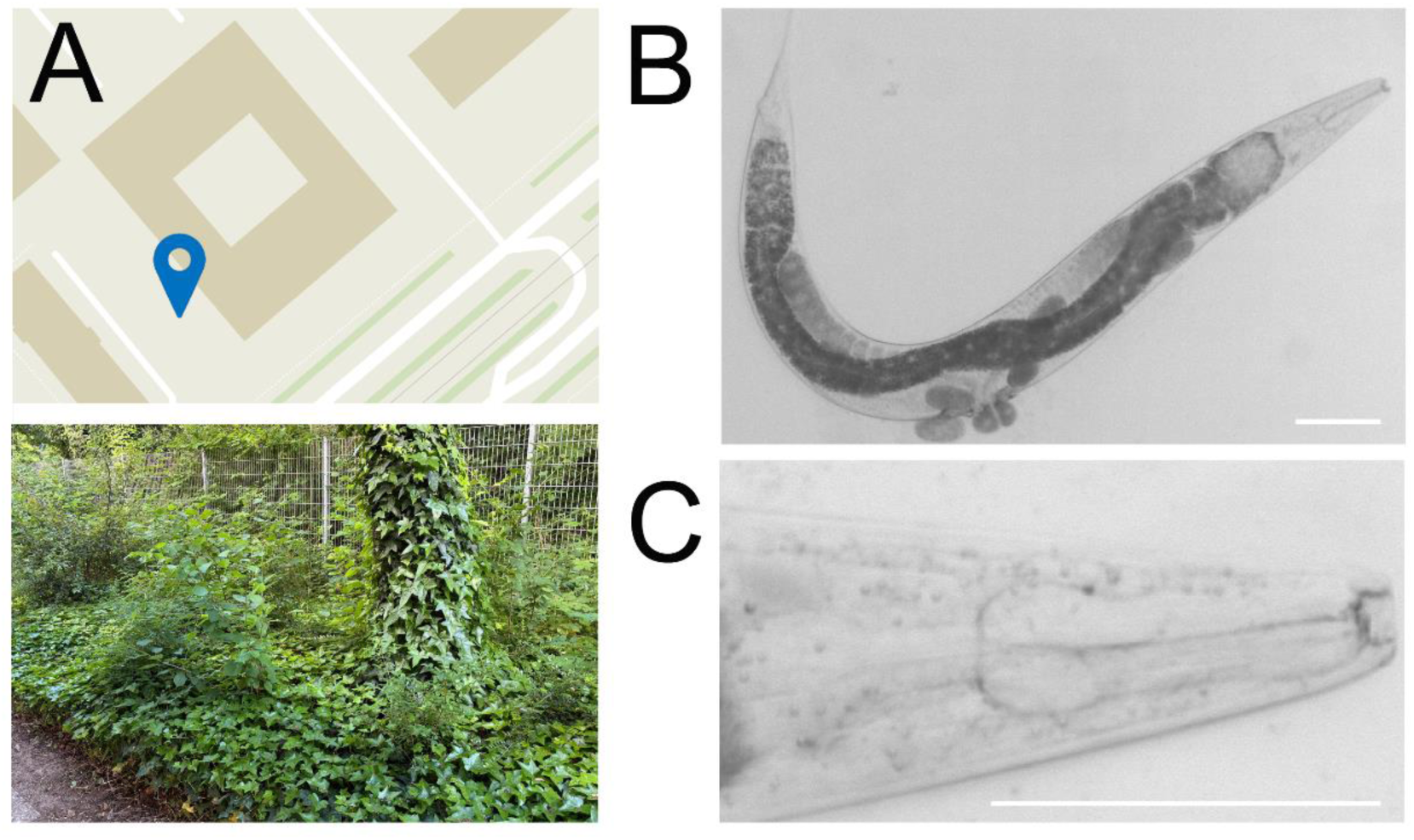
Isolation of diplogastrid nematodes from the field. (A) Schematic map of the field-isolation site located near a busy traffic road in Duesseldorf, Germany (coordinates: 51°12’19.2"N 6°47’10.6"E), and corresponding image of the sampling site. (B) Representative micrograph of an isolated nematode belonging to the Diplogastridae family. (C) Representative high-magnification micrograph shows the pharynx with the prominent teeth (right, dark gray) located at the base of a short stoma. Bars, 100 µm.

**Supplementary Figure 3.**
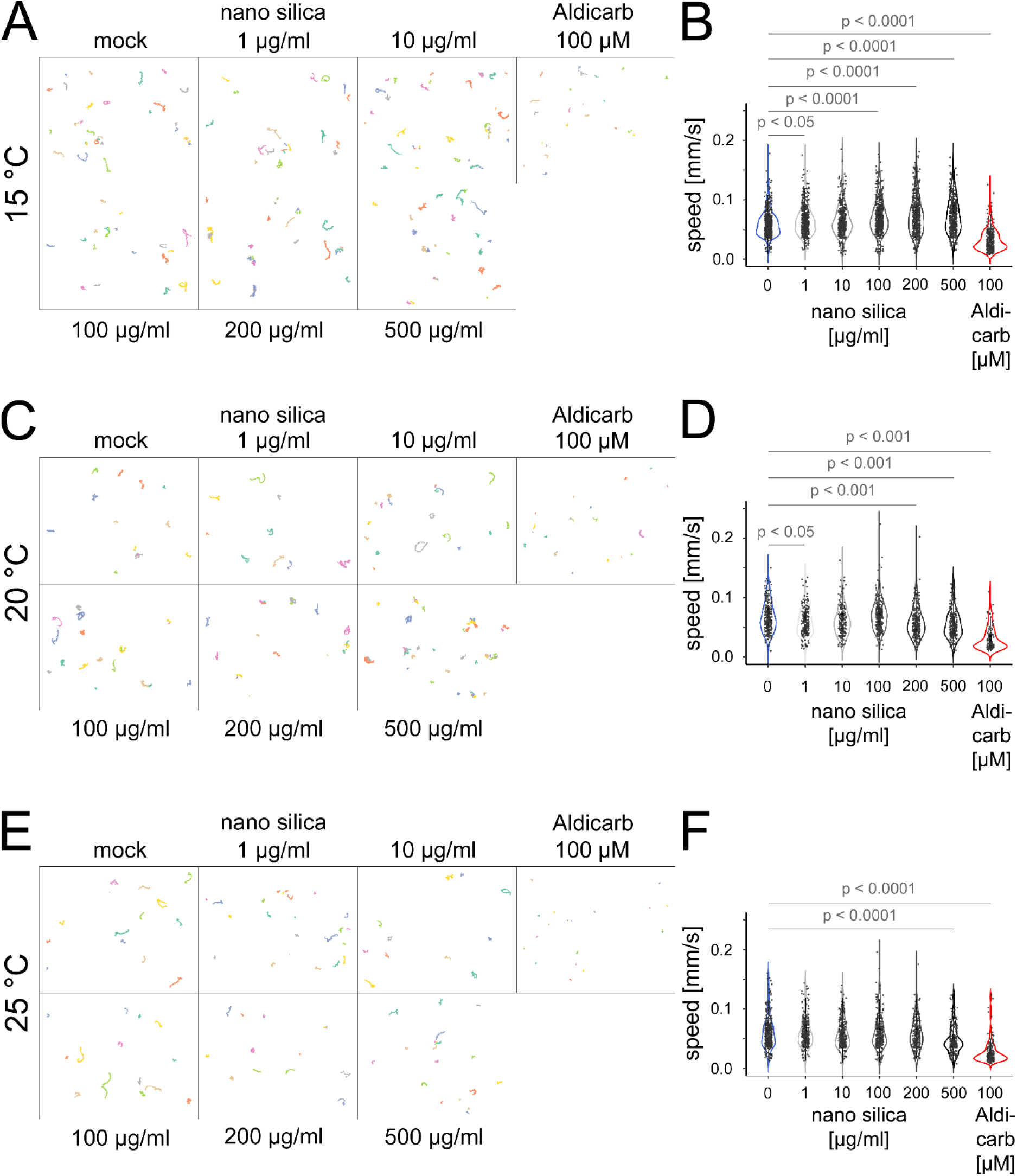
Combined effects of silica NPs and ambient temperature on field isolated rhabditid nematode locomotion. Representative swimming tracks of individual adult rhabditid nematodes (multi-colored) after exposure to silica nanoparticles (1-500 µg/mL) or Aldicarb (100 µM) at (A) 15 °C, (C) 20 °C, and (E) 25 °C. Short tracks indicate impaired locomotion, while longer tracks represent enhanced swimming. (B, D, F) Box plots display the average swimming speed of rhabditid field isolated nematodes at 15 °C (B), 20 °C (D), and 25 °C (F). Swimming activity is enhanced with increasing nanoparticle concentrations at 15 °C (A, B), whereas higher temperatures or the exposure to Aldicarb lead to reduced swimming locomotion (C-F). Data represent means ± SD from at least three independent experiments, with n ≥ 35 worms per condition in each experiment (Kruskal-Wallis ANOVA with Dunn’s post hoc test).

**Supplementary Figure 4.**
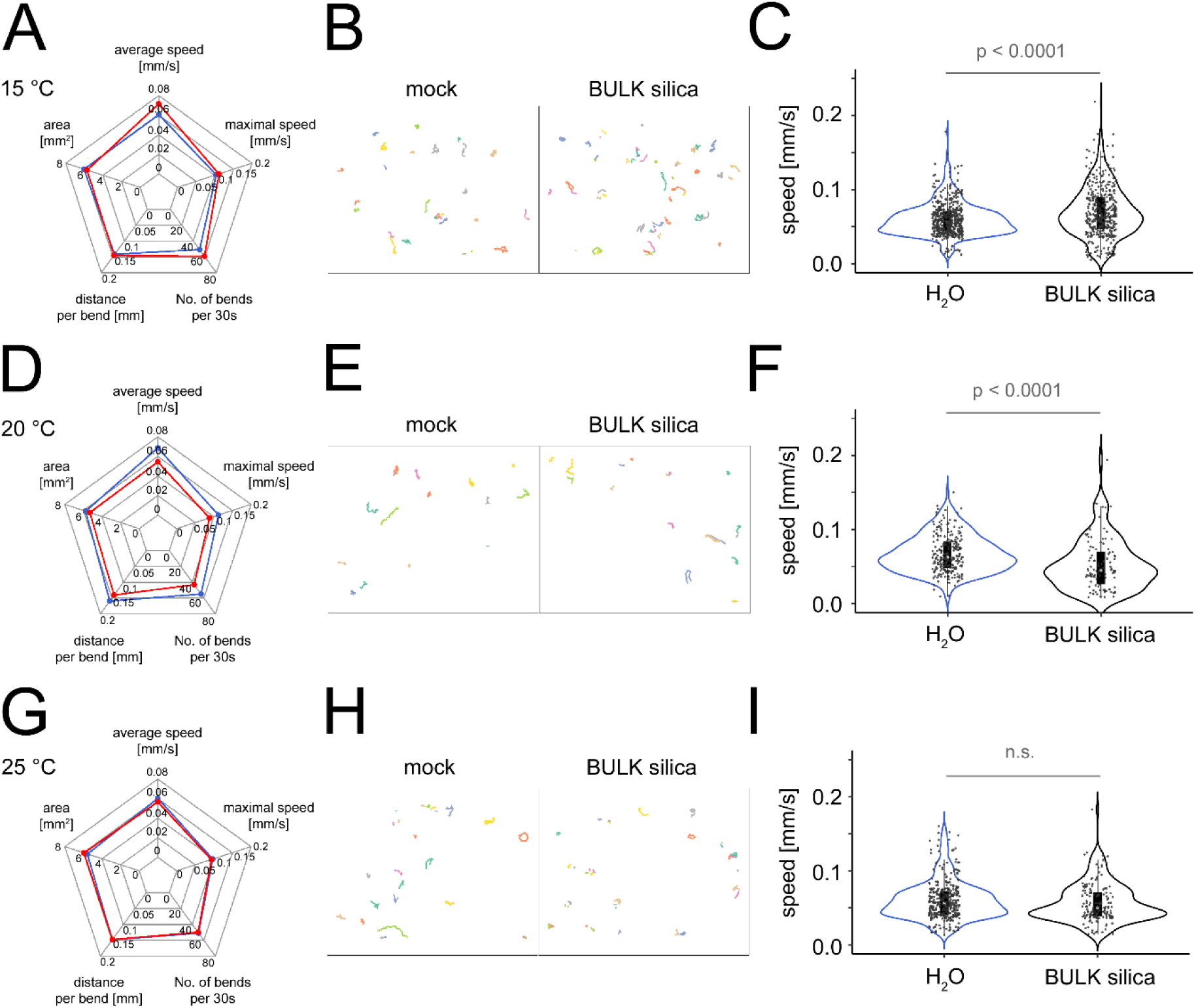
Effects of bulk silica particles and ambient temperature on field isolated rhabditid nematode locomotion. Adult field isolated rhabditid nematodes were exposed for 72 h to 500 µg/ mL BULK silica or H_2_O (control) at 15 °C (A-C), 20 °C (D-F) or 25 °C (G-I). (A, D, G) Spider plots show mean values of key locomotion parameters: average speed, maximum speed, number of bends per 30 seconds, distance per bend, and area. Control data are shown in blue and BULK silica–exposed groups are displayed in red. (B, E, H) Representative swimming tracks (30 seconds) of individual worms (multi-colored) are shown for control and BULK silica conditions. Short tracks indicate impaired locomotion, while longer tracks represent enhanced swimming. (C, F, I) Quantification of average speed in mock control and BULK silica-treated worms. At 15 °C, swimming activity increased upon exposure (A-C), while at 20 °C, it declined (D-F). At 25 °C, no significant change was observed (G-I), particularly in average speed and number of bends, indicating these as the most sensitive locomotion parameters to BULK silica exposure. Data represent mean ± SD from three independent experiments, with n ≥ 37 worms per condition in each experiment. Statistical analysis was performed using Kruskal–Wallis ANOVA followed by Dunn’s post hoc test.

**Supplementary Table 1.** Single-worm tracking of *C. elegans* and rhabditid field isolates.

| Category | Parameter | Description | *C. elegans* | Wild Rhabditid Isolates |
| --- | --- | --- | --- | --- |
| Worm Throughput | Total worms analyzed | Number of individual worms tracked | 11,029 | 21,486 |
| Worm Throughput | Replicates per condition | Biological replicates (each including technical repl.) | 21-146 worms × 4 replicates | 25–166 worms × 3–7 replicates |
| Exposome Factors | Nanoparticles (NPs) tested | Types of silica NPs | Monodisperse NPs, Aerosil 90, Aerosil 200 | Same |
| Exposome Factors | Nanoparticle concentrations | Silica NPs dose range | 1, 10, 100, 200, 500 µg/mL | Same |
| Exposome Factors | Temperatures tested | Environmental test conditions | 15°C, 20°C, 25°C | Same |
| Exposome Factors | Aldicarb | Positive control neurotoxicant | — | 100 µM for 30 min |
| Experimental Design | Exposure duration | Duration of NP exposure | 72 hours | Same |
| Experimental Design | Measured behavior | Locomotion metrics recorded | Speed, # bends, distance/bend, area | Same |
| Data Output | Tracking resolution | Image capture rate and duration per worm | 21 fps for 30 seconds | Same |
| Data Output | Quantitative features | Data features extracted | Speed, # bends, distance/bend, area | Same |
| Data Output | Total data points | Total individual measurements | 55,145 | 107,430 |

## Notes

### Competing Interest Statement

The authors have declared no competing interest.

